# Communication dynamics in the human connectome shape the cortex-wide propagation of direct electrical stimulation

**DOI:** 10.1101/2022.07.05.498875

**Authors:** Caio Seguin, Maciej Jedynak, Olivier David, Sina Mansour L, Olaf Sporns, Andrew Zalesky

## Abstract

Communication between gray matter regions underpins all facets of brain function. To date, progress in understanding large-scale neural communication has been hampered by the inability of current neuroimaging techniques to track signaling at whole-brain, high-spatiotemporal resolution. Here, we use 2.77 million intracranial EEG recordings, acquired following 29,055 single-pulse electrical stimulations in a total of 550 individuals, to study inter-areal communication in the human brain. We found that network communication models—computed on structural connectivity inferred from diffusion MRI—can explain the propagation of direct, focal electrical stimulation through white matter, measured at millisecond time scales. Building on this finding, we show that a parsimonious statistical model comprising structural, functional and spatial factors can accurately and robustly predict cortex-wide effects of brain stimulation (out-of-sample *R*^2^=54%). Our work contributes towards the biological validation of concepts in network neuroscience and provides insight into how white matter connectivity shapes inter-areal signaling. We anticipate that our findings will have implications for research on macroscale neural information processing and the design of brain stimulation paradigms.

---

Communication between neural elements is paramount to brain function. The complex network of gray matter regions interlinked by white matter fibers—the structural connectome—provides the anatomical substrate for macroscale communication in the brain [1, 2]. Communication via white matter connections underpins interactions between gray matter regions, facilitating the synchronization of neural activity (i.e., functional connectivity) along multiple spatiotemporal scales and giving rise to the brain’s rich functional dynamics [3–5]. Communication processes therefore constitute a bridge between structural and functional descriptions of nervous systems [6, 7].

It is well established that the functional connectivity between two anatomically connected regions is correlated to their structural connectivity [8–10]. Less well understood, however, are observations of co-activation between anatomically *unconnected* regions. For example, functionally coupled areas within the same intrinsic resting-state network do not always share a direct structural connection [11], as is the case for parts of the parietal cortex and the precuneus involved in the default mode network [12, 13]. Functional connectivity has also been observed between homotopic gray matter loci—normally connected by inter-hemispheric callosal fibers—following complete severance of the corpus callosum [14, 15]. Similarly, targeted pharmacogenetic deactivation of single gray matter structures has been reported to cause global changes in inter-areal co-activation patterns, impacting the activity of regions that were not anatomically connected to the manipulated site [16]. These examples illustrate that large-scale brain function is not supported exclusively by interactions between directly connected regions, but also by communication dynamics that facilitate polysynaptic information integration between distant and anatomically unconnected regions [7].

Numerous network communication models have been proposed to describe polysynaptic signal propagation in the connectome [17]. Computed on structural connectivity, these models quantify the ease of communication between pairs of regions by considering a putative strategy to guide signal transmission. Network communication models provide tractable interpretations of the interplay between structural connectivity and inter-areal interactions, and can thus be used to form and test hypotheses about the relationship between brain structure and function. An emerging body of evidence indicates that these models can explain inter-individual variation in cognitive [18, 19] and clinical [20–24] variables, as well as various aspects of functional and effective connectivity derived from blood-level-oxygen dependent (BOLD) fMRI time courses [16, 25–34].

Despite this progress, direct evidence that network communication models reflect aspects of biological neural signaling remains lacking. Slow-fluctuating BOLD time series—the focus of the vast majority of studies to date—are unable to capture inter-areal communication the at millisecond time scales inherent to signal propagation. Conversely, high-temporal-resolution electrophysiological recordings are typically limited in their ability to measure spatially-resolved neural activity across the whole brain [35, 36]. These technological obstacles have thus far hampered the biological validation of brain network communication models.

Here, we address this challenge using invasive stereotactic electroencephalography (SEEG) recordings of cortico-cortical evoked potentials (CCEPs) following direct electrical stimulation of the human brain. We use data from the F-TRACT project [37–39] to derive whole-brain, high-spatial-resolution maps of stimulus propagation measured at millisecond resolution. We hypothesize that modeling network communication on the human structural connectome can predict the cortex-wide propagation of focal electrical stimulation. Importantly, we consider a wide range of models—from efficient routing protocols to passive diffusion processes—to determine which conceptualizations of network communication best explain stimulus transmission. In parallel, we leverage the interpretability of our modeling framework to make inferences about which organizational properties of the connectome contribute to inter-areal communication. We posit that progress in understanding the spread of exogenous, interventional stimulation will provide insights into the underlying mechanisms governing large-scale neural communication.

## RESULTS

The F-TRACT project [37, 38] processed multicenter SEEG data acquired in a large sample of patients with drug-resistant epilepsy. The position and number of implanted electrodes were separately determined for each patient based on prior clinical knowledge. Patients underwent focal single-pulse stimulation and invasive recordings of neural activity at millisecond resolution. Compiling data from a total of 550 participants, 29,055 stimulations and 2.77 million pairs of recording electrodes, allowed the computation of whole-brain, group-level probability (P) and amplitude (A) matrices of regional activation following stimulation (Fig 1a; *Materials and Methods*). The matrix entries *P_ij_* ∈ [0,1] and 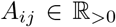 capture, respectively, the probability of observing a significant excitatory response in region *j* following stimulation of region *i*, and the median amplitude of significant responses.

**FIG. 1.**
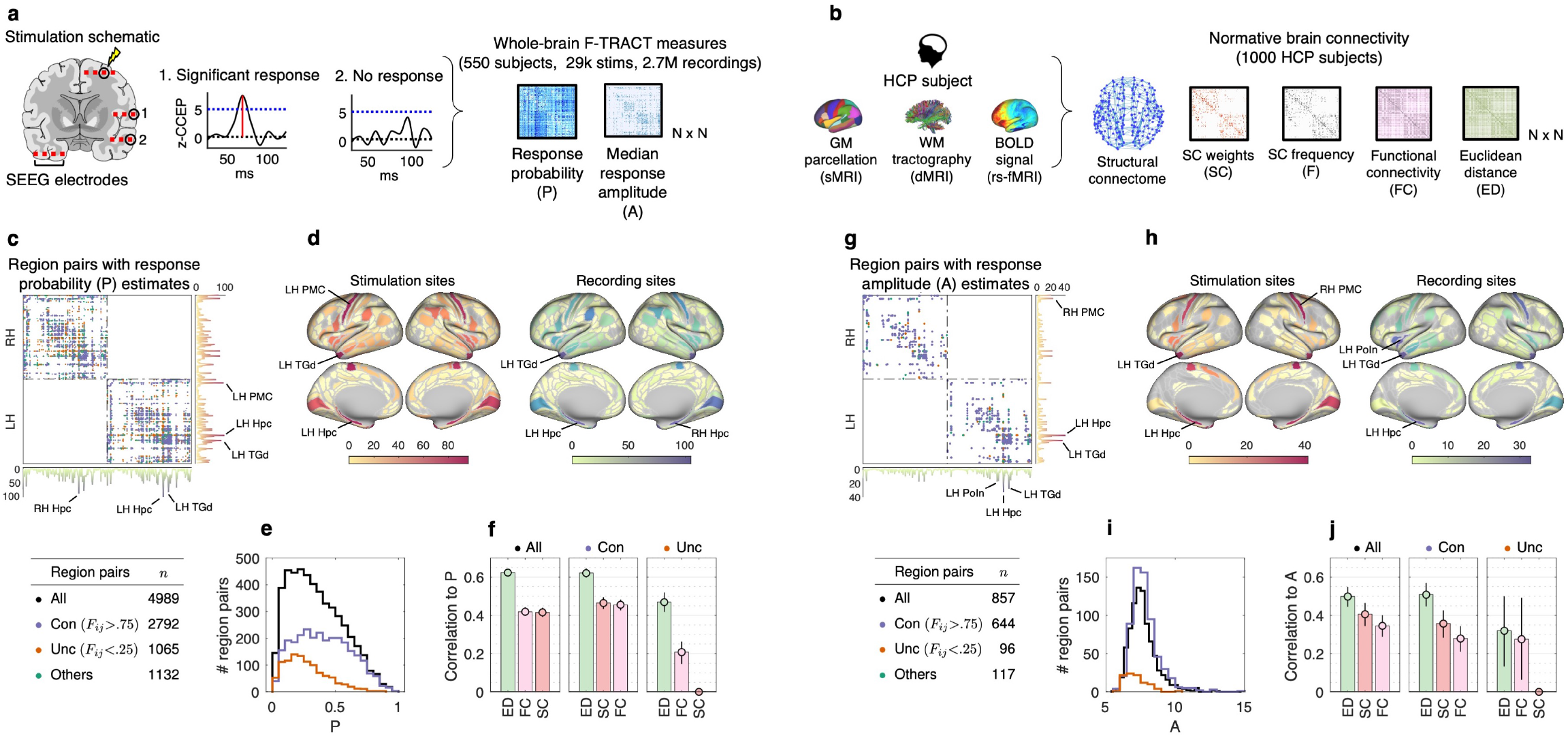
**(a)** Schematics of the F-TRACT project. Subjects were implanted with SEEG electrodes capable of delivering localized stimuli and recording electrical activity at millisecond resolution. The toy example shows a stimulation performed on a single subject, where the effects of a single-pulse stimulus are recorded (schematics are illustrative and do not reflect the bipolar stimulation montage and the true spacing between contacts). A significant response is measured by contact 1, as defined by a *z*-scored CCEP larger than 5. Contact 2 does not record a significant response. Based on its spatial coordinate, each contact was assigned to a cortical gray matter region. Whole-brain matrices of response probability (P) and median amplitude (A) were computed by combining data from 29,055 stimulations in 550 subjects. **(b)** Normative, group-representative measures of brain connectivity were inferred from 1,000 healthy participants of the Human Connectome Project using structural, diffusion, and resting-state functional MRI data. **(c)** Pairs of regions used to model response probability. Violet and copper matrix entries denote regions that are likely and unlikely to be anatomically connected, respectively. Warm- and cool-colored line plots indicate the number of times each region featured as a stimulation and recording site, respectively, and were projected onto the cortical surface on panel **(d)**. **(e)** Response probability distributions for all (black), connected (violet), and unconnected (copper) region pairs. **(f)** Bars show the absolute value of Spearman’s *ρ* between response probability and ED, SC and FC, computed for all (left), connected (center), and unconnected (right) region pairs. Circles and error bars indicate the mean and 95% confidence interval correlations obtained from 1,000 bootstrapped samples. **(g–j)** Same as (c–f) but for response amplitude.

To investigate the role of brain connectivity on stimulus response, we mapped structural and functional brain networks from MRI data of 1,000 healthy participants of the Human Connectome Project [40] (HCP; Fig 1b; *Materials and Methods*). We generated four group-representative normative measures: structural connectivity (SC), structural connection frequency (F) across HCP participants, functional connectivity (FC) and interregional Euclidean distance (ED). While our primary goal is to understand communication via white matter connectivity, considering functional and spatial factors enabled a comprehensive examination of how stimulus propagation is associated with multiple aspects of brain organization.

### Whole-brain maps of electrical stimulus propagation

We analyzed response probability estimates between 4,989 region pairs (15.4% of all intra-hemispheric region pairs; Fig 1c), out of which 2,792 were likely to be anatomically connected (*F* > 75%; violet matrix entries) and 1,065 were likely to be anatomically unconnected (*F* < 25%; copper matrix entries). Stimulation and recording sites covered the majority of the cortical surface (Fig 1d; 81% and 84% of regional coverage, respectively), and were primarily concentrated in areas involved in refractory focal epilepsy. For amplitude, we analyzed estimates between 857 region pairs (2.6% of all intra-hemispheric region pairs; Fig 1g), out of which 644 and 96 were likely and unlikely to be anatomically connected. Stimulation and recording sites remained distributed across large portions of the cortex (Fig 1h; 47% and 54% of regional coverage, respectively). Note that amplitude measurements are sparser since they only encode stimulations that elicited significant responses. Despite the heterogeneous spatial distribution of electrode implantations, pairwise response measures were distributed across multiple regions and systems, thus allowing for whole-brain analyses of stimulus propagation at high spatiotemporal resolution.

We found that anatomically connected regions showed higher response probability (Wilcoxon rank sum test *p* < 10^−68^; Fig 1e) and amplitude (*p* < 10^−12^; Fig 1i), compared to unconnected ones. As shown in Figs 1f,j, probability and amplitude were correlated to the SC (e.g., *ρ_con_* = 0.46,0.36 for P and A, respectively), FC (e.g., *ρ_all_* = 0.42, 0.34), and especially the ED (e.g., *ρ_all_* = −0.62, −0.50) between stimulation and recording sites (all *p* < 10^−32^).

In line with previous literature [41], these results indicate that multiple aspects of connectome architecture may shape stimulus propagation, including anatomical connectivity [10, 42], resting-state functional coupling [43, 44], and brain geometry [38].

### Modeling brain network communication

We next sought to investigate the network communication dynamics that facilitate polysynaptic (i.e., multi-step) stimulus propagation (Fig 2a). A network communication model determines a policy to guide signals through structural connectivity, and can therefore be used to quantify properties of stimulus transmission between regions. We considered five popular measures of brain network communication: shortest path efficiency [45, 46], navigation efficiency [47, 48], search information [25, 49], communicability [50, 51], and diffusion efficiency [52] (see Fig 2b for brief descriptions of measures and the *Materials and Methods* for technical details). Covering a wide range of signaling conceptualizations, these measures are positioned along a spectrum from routing—communication via efficient, selectively accessed paths—to diffusion—spreading along multiple network fronts [53].

**FIG. 2.**
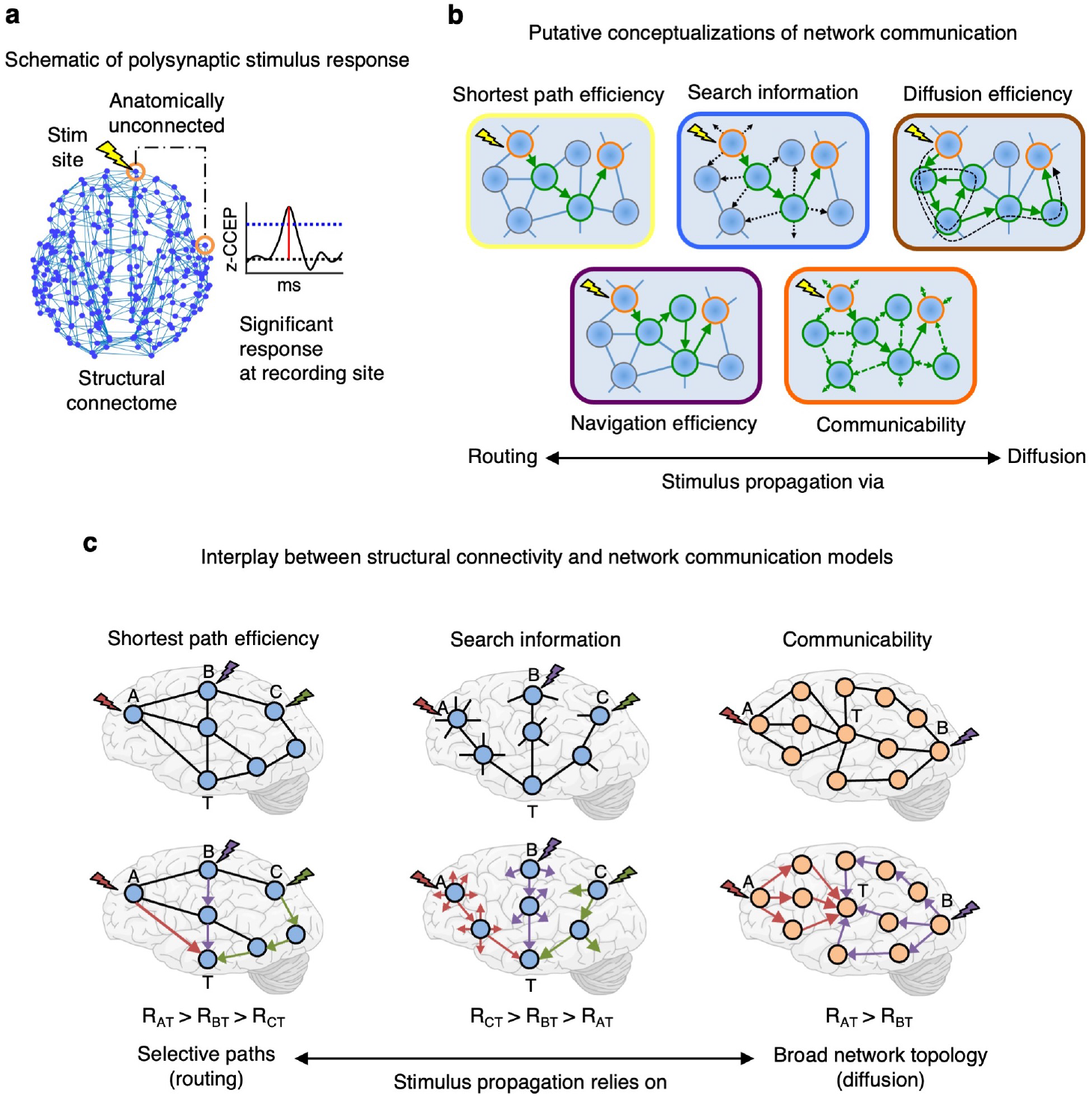
**(a)** A significant response is observed between anatomically unconnected stimulation and recording sites. How does white matter connectivity facilitate and constrain stimulus propagation? **(b)** Schematics of five putative measures of brain network communication. Briefly, shortest path efficiency (SPE) quantifies communication via the least costly path between regions; navigation efficiency (NE) presumes propagation via geometrically greedy paths; search information (SI) assumes a diffusive communication policy and measures the opportunity for signal dispersion along shortest paths; communicability (CMY) is a diffusive broadcasting process; and diffusion efficiency (DE) models propagation via random walks. **(c)** Network communication measures rely on different aspects of structural connectivity to quantify inter-areal propagation. Left: SPE models stimulus response (*R*) between stimulation (*A, B*, *C*) and recording (*T*) sites based exclusively on properties of the shortest—e.g., fewest intermediate regions—path between them. Center: SI takes into account the local connectivity along shortest paths. A stimulus propagated via regions with many connections is more prone to dispersion, leading to weaker responses. Despite being connected to *T* via a single intermediate region, *A*, *B*, and *C*, in this order, provide decreasing opportunities for signal dispersion. Right: Communicability is a broadcasting process that unfolds through the entire network. Short paths contribute more to communication than long ones. Despite both being connected to *T* via a single intermediate region, stimulation to *A* evokes stronger responses than stimulation to *B*, since *A* has a larger number of short paths to *T*.

Importantly, models located at different positions of the routing–diffusion spectrum rely on distinct properties of the underlying structural connectivity to guide signaling (Fig 2c). Routing measures (shortest path efficiency and navigation efficiency) consider that stimuli are communicated exclusively via efficient paths comprising a small number of high-weight connections. In contrast, diffusion measures (communicability and diffusion efficiency) posit that the broader network topology also plays a significant role in communication. For example, the presence of multiple alternative paths from stimulation to recording sites is expected to increase the strength of stimulus response. Lastly, search information combines aspects of both routing and diffusion. In the context of brain stimulation, it can be interpreted as the propensity for a stimulus to be dispersed and diluted while traveling via the shortest path from stimulation to recording sites.

Determining whether certain measures yield better explanations of stimulus response than others can therefore provide insight into which topological properties of the connectome are most relevant for neural communication.

### Network communication models explain polysynaptic propagation of electrical stimulation

We correlated network communication measures computed on the structural connectome with response probability and amplitude. To account for the contribution of spatial proximity between stimulation and recording sites, we computed both full (*ρ*) and partial (*ρ*′) correlations, the latter controlling for the effect of ED.

We first observed that, despite the strong influence of ED, connectivity-based measures explained significant variance in stimulus propagation that was not accounted for by spatial proximity (Fig 3c–h; dark-colored bars). More specifically, we found significant partial correlations between probability and network communication measures (e.g., search information: 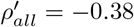; Fig 3c, blue bar), SC (e.g., 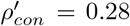; Fig 3d, red bar), and FC (e.g., 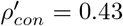, Fig 3d, pink bar), with comparable findings observed for response amplitude (Fig 3f–h).

**FIG. 3.**
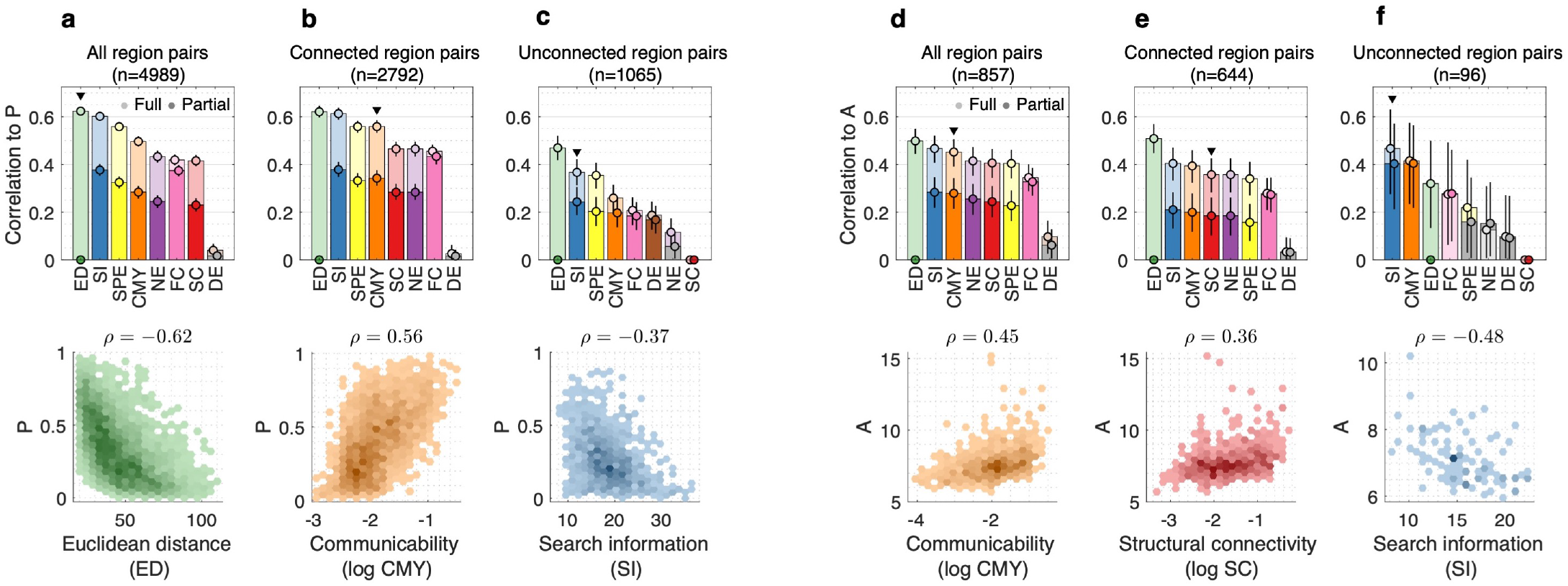
**(a-b)** Spearman’s correlations to response probability for (a) all, (b) anatomically connected and (c) anatomically unconnected region pairs. Absolute values of full and partial (controlling for ED) correlations are shown in light- and dark-colored bars, respectively. Gray bars indicate correlations that were not significant at *α* = 5%. Circles and error bars indicate, respectively, the mean and 95% confidence interval correlations obtained from 1,000 bootstrapped samples. Black triangles mark full correlations illustrated by scatter plots in the lower panels. Each data point in the scatter plots represents a pair of regions, with darker colors indicating higher point density. **(d-f)** Same as (a-b) but for response amplitude.

Models of network communication explained significantly greater variability in stimulus propagation than structural connectivity alone (Fig S1). In particular, search information consistently ranked as the most explanatory network measure of response probability and amplitude (e.g., *ρ_all_* = −0.60, −0.47, Figs 2c,h, respectively; both *p* < 10^−32^), in many cases yielding statistically greater correlations than alternative measures (Fig S1). Interestingly, top-performing network measures, such as search information and communicability, surpassed the explanatory power of SC weights even for associations computed exclusively for pairs of anatomically connected regions (Fig S1b,e). This suggests that stimulus transmission between connected sites may not be supported entirely by direct fibers, but also facilitated by propagation through intermediate regions and connections in the surrounding network topology.

Response amplitude measurements between anatomically unconnected regions describe instances of significant multi-step stimulus transmission through the connectome, thus reflecting a primary conceptual application of network communication models (Fig 2a,b). Crucially, in this scenario (Fig 3f), search information and communicability markedly outperformed alternative network measures and ED (SI: *ρ_unc_* = −0.48, *p* = 8 × 10^−7^, CMY: *ρ_une_* = 0.42, *p* = 2 × 10^−5^; Fig S1f,l). These results provide evidence that network communication measures can explain the propagation of electrical stimuli—traveling at millisecond resolution—between anatomically unconnected regions.

### Multivariate prediction of electrical stimulus propagation

Next, we combined SC, FC, ED, and all network communication measures into a single multivariate predictive model (*Materials and Methods*). Using a machine learning method that penalizes model complexity, we sought to identify sparse sets of predictors that explained non-overlapping variance in response probability and amplitude. We found that a simple predictive model comprising Euclidean distance (40.2% average contribution), search information (32.5%) and FC (24.6%) explained *R*^2^ = 54% of out-of-sample variance in response probability (Fig 4a–c). For amplitude, a combination of Euclidean distance (56.5% average contribution), communicability (29.5%), as well as search information and FC (14% joint contribution), yielded out-of-sample *R*^2^ = 31% (Fig 4d-f).

**FIG. 4.**
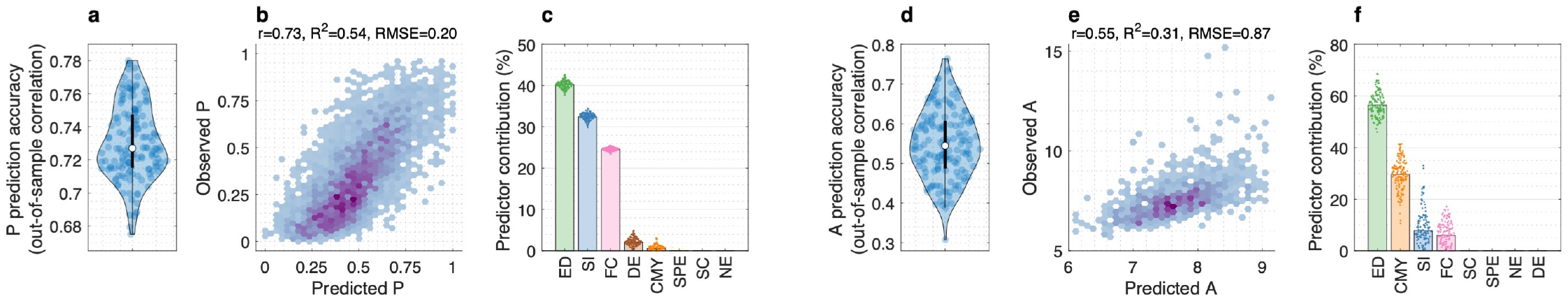
Machine learning predictions of stimulus propagation. A lasso regression model was trained to predict stimulus response from a combination of SC, FC, ED, and all network communication measures. By penalizing model complexity (number of predictors), the lasso regression seeks to identify a sparse set of input variables that accurately predicts the outcome variable. **(a)** Out-of-sample prediction accuracy for 15 repetitions of 10-fold cross validation, computed as the Pearson correlation between predicted and observed probabilities. Scatter plot of predicted versus observed probabilities with prediction accuracy evaluated as the Pearson correlation, *R*^2^, and root mean squared error (RMSE). **(c)** Average contribution of input predictors across 15 repetitions of 10-fold cross validation. **(d–f)** Same as (a–c) but for predictions of response amplitude.

These results corroborate and complement our univariate analyses, indicating that a parsimonious combination of structural, functional and spatial factors can accurately predict both the likelihood and the strength of cortex-wide responses to focal brain stimulation. Additionally, these analyses further indicate that search information and communicability provide the most explanatory conceptualizations of stimulus propagation through structural connectivity.

### Control analyses

We performed multiple control analyses (*Materials and Methods*) and found that our results remained consistent across a number of methodological choices (Fig S2), including replication in an independently acquired and processed normative connectivity dataset (Figs S3, S4). In addition, using ensembles of randomized null networks, we observed that our findings were contingent on the empirical topology of SC, such that rewiring of connections markedly reduced correlations between network communication measures and stimulus response (Fig S5).

## DISCUSSION

Here, we used a large dataset of direct electrical stimulation to study inter-areal communication in the human brain. Considering the causal effects of interventional stimulation allowed us to characterize the propagation of electrical pulses without the need for statistical or information-theoretical measures of information transfer between regions. In addition, we leveraged intra- and inter-subject variability in electrode placement to derive whole-brain, high-spatiotemporal-resolution maps of stimulus propagation. Together, these factors enabled us to test whether computational modeling of communication dynamics in the human connectome can explain empirical stimulus transmission through the brain.

### Towards the biological validation of brain network communication models

Taken together, our analyses identified two key findings. First, we provide evidence that network communication models—computed on structural connectivity inferred from tractography and MRI—reflect patterns of spatially-resolved, millisecond resolution, stimulus propagation measured across the whole brain. Our results recapitulate previous findings that structural connectivity shapes fast brain dynamics [10], and corroborate the use of network communication models to study neural information processing in healthy [54, 55] and clinical populations [20–24].

Second, adding to previous literature [16, 25, 41, 50], we found that—across multiple control analyses—search information and communicability led to the most accurate and robust accounts of stimulus propagation. This included correlations to the response amplitude between anatomically unconnected regions, indicating that these measures can explain multi-step propagation through the connectome. Despite being computed on the same structural substrate, alternative measures yielded weaker or inconsistent associations, and did not provide additional explanatory power when considered in multivariate statistical models. This specificity in predictive utility constitutes an important step towards determining which network communication models are more suitable to describe biological neural signaling.

Our work complements parallel research on network control theory [56, 57] and biophysical models of neural activity [58–60]. Recent reports have highlighted similarities and opportunities for synergy between these approaches for modeling brain function [61, 62]. As such, the present evidence in support of network communication models also bolsters research on these alternative directions. In particular, our work calls for the extension of efforts towards the empirical and biological validation of computational models of macroscale brain function.

### Understanding how brain network organization facilitates inter-areal stimulus propagation

A key feature of network communication models is that they provide tractable interpretations of the interplay between brain structure and function. Our findings reveal insights into how topological properties of structural connectivity facilitate inter-areal signaling (Fig 2). Search information and communicability conceptualize network communication as a diffusive process, in which stimulus propagation is shaped by signal dispersion and transmission via multiple network fronts. This notion challenges the mainstay assumption in network neuroscience that communication occurs exclusively via shortest paths [48, 52], and suggests that polysynaptic dynamics may play a role even for anatomically connected regions.

We envision that these results may contribute to the design of brain stimulation paradigms that—beyond maps of direct connectivity—also leverage knowledge of network communication dynamics [63]. For example, models of polysynaptic transmission could inform the selection of stimulation sites for indirect modulation of deep gray matter structures by non-invasive techniques, such as transcranial magnetic stimulation [64, 65].

In addition to propagation constrained by white matter, spatial and functional factors were also predictive of stimulus response. We note that while the influence of spatial proximity may reflect neurophysiological evidence that amplitude decays as a function of axonal length [38], it could also be confounded by factors such as non-axonal conduction, spatial autocorrelations in MRI data [66], and inaccurate reconstructions of short-range fibers by tractography [67]. We also found that, despite a relatively weaker explanatory power, functional connectivity explained unique variance in stimulus propagation. In line with previous studies [43, 44], this indicates that intrinsic functional dynamics may relate to stimulus response in ways complementary to structural and spatial factors [60]. Multimodal models of connectome communication that incorporate information on both structural and functional connectivity may therefore provide a fruitful direction for future research [68, 69].

### Limitations and future directions

A number of limitations of the presence work should be discussed. First, stimulus response were measured in individuals with refractory epilepsy. While conservative methods were applied to filter out pathological activity from our analyses (see *Materials and Methods*), the extent to which recordings from epilepsy patients are representative of healthy brain activity remains an active topic of research [36, 39]. Similarly, CCEPs elicited by direct electrical stimulation may not reflect physiological patterns of inter-areal communication. While future work is required to investigate these issues, we note that our findings are congruent with previous reports that diffusion-based communication models explained activity measured using electrocorticography in the absence of electrical stimulation [41] and non-invasive magnetoencephalography in healthy adults [70].

Our analyses focused on intra-hemispheric stimulus propagation (Fig. 1c,g). This was a result of selecting region pairs for which stimulus response measures were reliably estimated based on a large number of stimulations (see *Materials and Methods*). Nonetheless, determining whether our results extend to inter-hemispheric communication is an important step for subsequent research. Lastly, while our results corroborate the use of group-level normative connectivity to study neuromodulation in clinical populations [71], future research on modeling subject-level data is necessary.

### Conclusion

In conclusion, through the analyses of more than 2.7 million SEEG recordings, we provide novel insights into connectome communication and polysynaptic stimulus propagation. This work is a step towards the development and validation of biologically realistic models of brain communication. Our results provide new avenues for research on macroscale neural information processing and brain stimulation paradigms that leverage network communication dynamics.

## MATERIALS AND METHODS

### F-TRACT dataset

The F-TRACT project is a continuing effort in collecting, processing and compiling iEEG recordings of drug-resistant epilepsy patients being prepared for resective surgery [37, 38]. In this study, we used data from 550 patients (50% women, age 31 ± 10 years old) collected across 20 epilepsy centers in Europe, North America and Asia (https://f-tract.eu/consortium). Patients were implanted with intracerebral depth electrodes (stereo-electroencephalography; SEEG) with an average of 87 ± 37 electrode contacts per participant. Cortico-cortical evoked potentials (CCEPs) were recorded following low-frequency, bipolar direct electrical stimulation with the following parameters: 1 Hz stimulation, biphasic (79% of CCEPs) or monophasic (21%) electrical pulses, 3.3 ± 1.2 mA mean pulse intensity, 1.0 ± 0.4 ms mean pulse duration, and 3.3 ± 1.5 *μ*C mean pulse charge.

CCEPs were processed as follows. First, SEEG signals were preprocessed to identify stimulation events, filter bad channels [72], and correct for stimulation artifacts [73]. Recordings with high likelihood of reflecting pathological activity were identified and discarded using Delphos software to mitigate the impact of epileptogenic processes in further analyses [39, 74]. CCEPs on each electrode contact were baseline corrected and *z*-scored based on the [−400, −10] ms time window before stimulation. A stimulus response was deemed significant if a *z*-CCEP above *z* = 5 was observed within 800 ms of the stimulation. The response amplitude of a single stimulation was defined as the first *z*-CCEP peak above the significance threshold.

The position of electrode contacts was computed based on a patient’s structural MRI (T1-weighted imaging) and computerized tomography scans. Contacts placed in gray matter were assigned to the corresponding regions of the HCP-MMP1.0 parcellation [75]. For contacts placed in white matter, we first computed a 3 mm radius sphere centered in the contact’s coordinates and assigned them to the gray matter parcel with highest voxel representation within the sphere.

Whole-brain, group-level response probability (P) and amplitude (A) matrices were computed by compiling CCEP data from 2,769,985 SEEG recordings following 29,055 electrical stimulations across 550 individuals. The matrix entries *P_ij_* ∈ [0,1] and 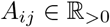 capture, respectively, the probability of observing a significant excitatory response in region *j* following stimulation of region *i*, and the median *z*-scored CCEP amplitude of significant responses. Note that P and A are asymmetric, such that *P_ij_* may differ to *P_ij_*. Data from the F-TRACT project is openly available at https://f-tract.eu/ and further details on data acquisition and processing are provided in [37–39].

### Normative measures of brain connectivity: HCP dataset

Structural connectivity (SC) was inferred from minimally preprocessed, high-resolution diffusion-weighted MRI data from 1,000 healthy young adults participating in the Human Connectome Project. Details of diffusion MRI acquisition and preprocessing are described in [76, 77]. Whole-brain white matter tractograms of individual participants were mapped using a probabilistic tractography pipeline (MRtrix3 software [78], multi-shell multi-tissue constrained spherical deconvolution [79], anatomically constrained tractography [80], iFOD2 tracking algorithm [81], 5M streamlines seeded from the gray-white matter interface; see [82] for further details). Cortical gray matter was divided into 360 regions using the HCP-MMP1.0 parcellation [75], while the Freesurfer parcellation [83] was used to define 14 subcortical structures (left and right thalamus, caudate, putamen, pallidum, hippocampus, amygdala and accumbens area). Note that while stimulus response measures were available only in the cortex, the inclusion of subcortical regions in structural connectomes enables the modeling of propagation mediated by the subcortex. The structural connectivity weight between two gray matter regions was defined as the number of streamlines between them divided by the sum of their surface areas, approximated as the number of voxels in the gray-white matter interface of each region [29]. Connections comprising fewer than 5 streamlines were pruned to reduce the high false positive rate of probabilistic tractography [84]. A group-representative connectivity matrix was computed using a consensus method that combined individual-level networks while preserving the average connection density across subjects [85]. This resulted in a final 374 × 374 SC matrix with 25% connection density.

Functional connectivity (FC) was mapped using minimally preprocessed, ICA-FIX resting-state functional MRI data from the same sample of 1,000 HCP participants. For each subject, four 14 minutes and 33 second scans (0.72 second TR) were acquired following details described in [77, 86]. Voxel-level blood-oxygen-level-dependent time series were linearly detrended, band-pass filtered and standardized [87]. Following previous work [88], a total of four nuisance variables were regressed out: the global signal (GS), the GS squared, the GS derivative, and the squared GS derivative. The time series of voxels located in the same gray matter parcel were averaged and FC was computed as the Pearson correlation between regional time series. A final 374 × 374 group-representative FC matrix was computed as the average of 4,000 individual matrices (4 sessions per subject for 1,000 subjects).

### Normative measures of brain connectivity: MICA dataset

The Multimodal Imaging and Connectome Analysis (MICA) dataset is an openly available resource (https://portal.conp.ca/) containing structural and functional brain networks for 50 healthy adults (23 women; 29.5 ± 5.6 years old) [89]. Brain networks were inferred using independent acquisition, preprocessing and connectivity mapping pipelines. We utilized the MICA brain networks comprising 374 regions, delineated according to the same cortical and subcortical parcellations used in the HCP dataset. To account for the nearly fully-connected nature of MICA structural networks (average connection density 84.1% across subjects) and prune potential false positives [84], we thresholded individual-level SC matrices by keeping only the top-25% strongest structural connections. Group-representative structural and functional networks were then derived following the same procedures described above for the HCP dataset.

### Network communication models

Let 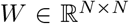 denote the matrix of SC weights between *N* regions, where *W_ij_* provides a measure of the strength and reliability of the anatomical connection between regions *i* and *j*. Anatomically connected and unconnected region pairs have *W_ij_* > 0 and *W_ij_* = 0, respectively. We define the matrix of SC lengths *L* = 1/*W*, where *L_ij_* quantifies the distance of travel cost between *i* and *j*. The remapping of connection weights into lengths is required for the computation of network communication models that aim to minimize the cost of transmitting signals between regions. Measures of network communication were computed using the Brain Connectivity Toolbox [90] and were defined as follows.

#### Shortest path efficiency

The Floyd-Warshal algorithm was used to identify the sequence of regions {*i, u*, …, *v, j*} such that 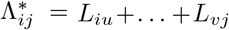 is the minimum transmission cost between regions *i* and *j*. Shortest path efficiency was defined as 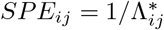 [45].

#### Navigation efficiency

Navigation is a greedy routing protocol that propagates signals based on a measure of inter-regional distance [47]. Following previous work [48], navigation from a source region *i* to a target region *j* was implemented as follows. First, identify *i*’s neighbor with the shortest Euclidean distance to *j* and progress to it. Repeat this process until *j* is reached (successful navigation) or a region is revisited (failed navigation). The length of successful navigation paths is given by Λ*_ij_* = *L_iu_* + … + *L_vj_*, where {*i*, *u*, …, *v, j*} is the sequence of nodes visited during routing, whereas failed navigation entails Λ*_ij_* = ∞. Navigation efficiency was defined as *N E_ij_* = 1/Λ*_ij_*.

#### Diffusion efficiency

The diffusion efficiency quantifies the ease of communication from region *i* to region *j* under the assumption that signals are transmitted via unbiased random walks [52]. In this scenario, the probability of traveling from *p* to *q* in one step is given by the transition matrix 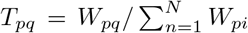. The transition probability matrix can be used to analytically compute 〈*H_ij_*〉, the average number of steps required for a random walker to travel from *i* to *j* [91]. Diffusion efficiency is defined as 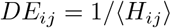.

#### Search information

Search information quantifies the amount of information necessary to bias a random walker to travel via shortest paths, capturing the accessibility of efficient routes under a diffusive model of communication [25, 49]. Considering the transition matrix *T* of a random walk process, *Pi_ij_* = *T_iu_*× … × *T_vj_* is the probability of a random walker serendipitously traveling from region *i* to region *j* via their shortest path {*i, u*, …, *v, j*}. Search information was defined as *SI_ij_* = –*log*_2_(Π_*ij*_).

#### Communicability

Communicability is defined as a weighted sum of the total number of walks between two regions, in which the contribution of each walk to signal propagation is inversely proportional to their length [51, 92]. Formally, 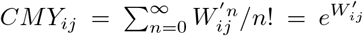, where 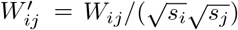 is a normalized connectivity matrix that attenuates the influence of high strength region and 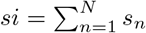 is the strength of region *i* [92].

### Univariate modeling of stimulus response and control analyses

SC, FC, ED, and the 5 network communication measures were correlated to response probability and amplitude using Spearman rank correlation. Associations were computed across pairs of cortical regions. In order to reduce potential effects of measurement noise and interindividual variability in CCEPs, we considered, in the main text, pairs of regions for which (i) a minimum of 100 stimulation experiments were available for the computation of probability, (ii) a minimum of 100 significant stimulation responses were available for the computation of amplitude, and (iii) SEEG recordings were acquired across a minimum of 5 different patients. In addition, to mitigate potential confounding factors relating to non-axonal volume conduction, spatial autocorrelations in MRI data [66], and inaccurate reconstructions of short-range U-fibers ny tractography [67], the results in the main text excluded pairs of regions closer than 20 mm in Euclidean distance [93].

We performed a number of control analyses to determine whether our findings were robust to variations in the methodological choices described above. Two sets of control analyses were considered. Firstly, in Fig S2, we systematically analyzed variations to the following parameters:

- Minimum number of stimulation experiments/significant responses between region pairs (100 [main text] vs. 50 [control analysis]).
- Minimum structural connectivity frequency across HCP subjects for considering two regions to be likely anatomically connected (75% [main text] vs. 90% [control analysis]).
- Maximum structural connectivity frequency across HCP subjects for considering two regions unlikely anatomically connected (25% [main text] vs. 10% [control analysis]).
- *z*-scored CCEP threshold for significant responses (*z* = 5 [main text] vs. *z* = 4, 6 [control analyses]).
- Minimum Euclidean distance between regions pairs (20 mm [main text] vs. 0 mm [control analyses]).

Secondly, in Figs S3, we re-examined all the scenarios above starting from structural and functional connectivity from the Multimodal Imaging and Connectome Analysis (MICA) dataset.

### Multivariate machine learning predictive model

Let 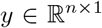 be a vector of outcome variables corresponding to the response probability or amplitude of *n* pairs of regions. Predictive modeling was carried out considering all pairs of regions used in the univariate analyses, so that *n* = 4989 for probability and *n* = 857 for amplitude. Let 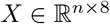 be a matrix of predictor variables comprising the SC, FC, ED, and the 5 measures of network communication between *n* pairs of regions. We used a lasso regression [94] to predict *y* from *X*.

The lasso regression was chosen as it implements a feature selection procedure aimed at identifying a sparse set of variables in *X* that accurately predicts *y*. This framework seeks to account for correlations between the variables in *X* by culling measures that offer redundant predictive utility. As such, predictor variables selected by the lasso regression tend to explain complementary portions of the variance in *y*.

Model training was done using 10-fold cross-validation by splitting *X* and *y* into train and test sets. Model fitting was performed exclusively on train sets and prediction accuracy was evaluated exclusively on test sets. To account for sensitivity to particular train-test splits, we repeated the 10-fold cross validation routine 15 times [95]. Computations were carried out using the cvglmnet MATLAB function [96].

More specifically, let {*X_a_, y_a_*} and {*X_e_, y_e_*} denote a split of *X* and *y* into train and test sets, respectively. The lasso regression computes 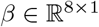 as

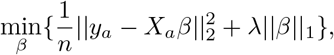

where the hyperparameter 0 ≤ λ ≤ 1 determines the sparsity of feature selection. The value of λ was tuned using nested cross-validation, in which each outer training set *X_a_* was further split into 10 inner training sets. For each inner training fold, we considered *K* logarithmically spaced λ from λ_*min*_ to *λ_max_*, where *λ_min_* = *ϵλ_max_* and *λ_max_* is the smallest parameter value such that ||*β_inner_*||_1_ = 0. Values of *K* and *ϵ* were dynamically determined by the nested cross-validation algorithm, with typical values around *K* = 100, and *ϵ* = 0.01 [97]. The mean squared error (MSE) of the resulting K models were then evaluated on the inner test folds. To promote sparsity, the largest λ with MSE within one standard error of the optimal MSE was selected [97]. The selected λ was then used to compute *β* for the training set *X_a_*. The prediction accuracy of each outer folder was evaluated as the Pearson correlation and the root mean squared error (RMSE) between 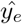 and *y_e_*, where 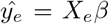. The contribution of predictor *i* was computed as 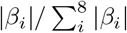. This procedure was repeated for 150 pairs of outer train and test sets (15 repetitions of 10-fold cross validation).

### Rewiring null models

SC topology was randomized using standard and cost-preserving versions of the Maslov-Sneppen rewiring algorithm [98]. Standard rewiring was performed by swapping pairs of connections while preserving empirical weight distribution and degree sequence. Cost-preserving rewiring was implemented by restricting swaps to pairs of connections with approximately the same length [99], as defined by the Euclidean distance between nodes. This routine preserves empirical weight distribution, degree sequence, connection length distribution, and connection weight-length relationship. In both routines, each connection was rewired once on average. Code implementing the standard and cost-preserving routines are available through references [90] and [99], respectively.

An ensemble of 1,000 standard and cost-preserving randomized structural networks was generated from empirical SC. Network communication measures were computed for each randomized network and correlated to response probability and amplitude to form distributions of null correlations (Fig S5). A network communication measure was considered to outperform a null model if its empirical correlation to stimulus response exceeded the 95th percentile of the null correlation distribution (non-parametric significance test at *α* = 5%). Evidence that a network communication measure outperforms a null model indicates that its association to stimulus response is not trivially explained by the SC attributes preserved during rewiring, and is instead contingent on the empirical topology of anatomical connectivity.

## ACKNOWLEDGMENTS

Brain imaging data were provided by the Human Connectome Project, WU-Minn Consortium (1U54MH091657; Principal Investigators: David Van Essen and Kamil Ugurbil) funded by the 16 National Institutes of Health (NIH) institutes and centers that support the NIH Blueprint for Neuroscience Research; and by the McDonnell Center for Systems Neuroscience at Washington University. CS was supported by the Australian Research Council (grant number DP170101815). MJ and OD were supported by funding from the European Research Council under the European Union’s Seventh Framework Programme (FP/2007-2013)/ERC Grant Agreement no. 616268 F-TRACT, the European Union’s Horizon 2020 Framework Programme for Research and Innovation under Specific Grant Agreement No. 785907 and 945539 (Human Brain Project SGA2 and SGA3), and from the Agence Nationale de la Recherche grant number ANR-21-NEUC-0005-01. SML is supported by a Melbourne Research Scholarship. AZ is supported by the National Health and Medical Research Council of Australia (APP1118153).

---

**FIG. S1.**
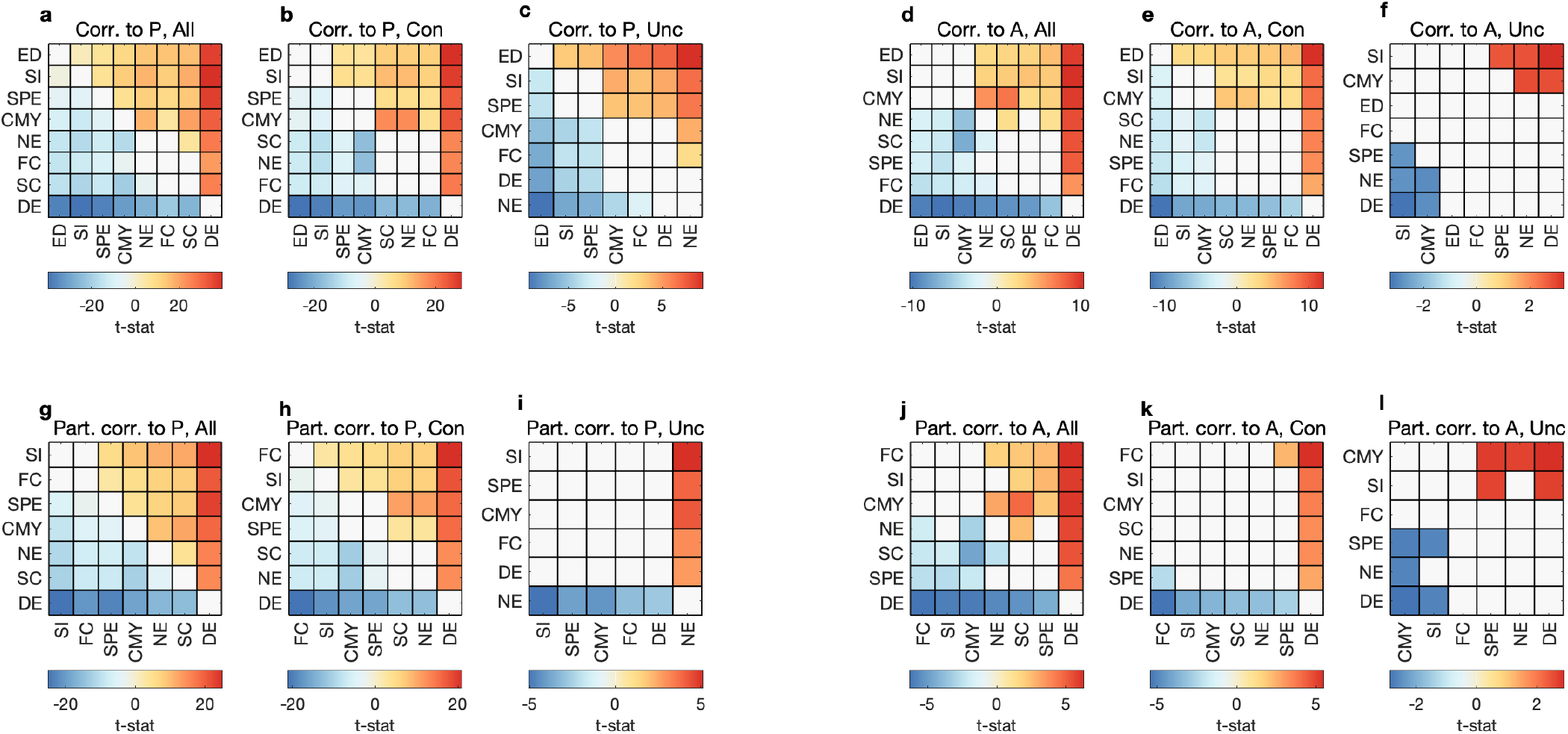
Pairwise *t*-statistics from two-sample *t*-tests performed on the distributions of 1,000 bootstrapped Spearman correlation coefficients presented in Fig 3. A warm-colored (cool-colored) matrix entry *ij* indicates that the mean correlation obtained for measure *i* is statistically higher (lower) than the mean correlation obtained for measure *j*. Matrices are symmetric about their main diagonal, with the opposite signs. A white matrix entry indicates no significant difference between the mean correlations obtained from two measures. Statistical tests for each *n × n t*-statistics matrix were adjusted using the Benjamini-Hochberg correction of the False Discovery Rate for (*n* × (*n* – 1))/2 multiple comparisons. Statistics are shown for full Spearman correlations with response probability for **(a)** all, **(b)** anatomically connected, and **(c)** anatomically unconnected region pairs. (d–e) Same as (a–c) but for response amplitude. (g–l) Same as (a–f) but for partial Spearman correlations controlling for the effect of Euclidean distance.

**FIG. S2.**
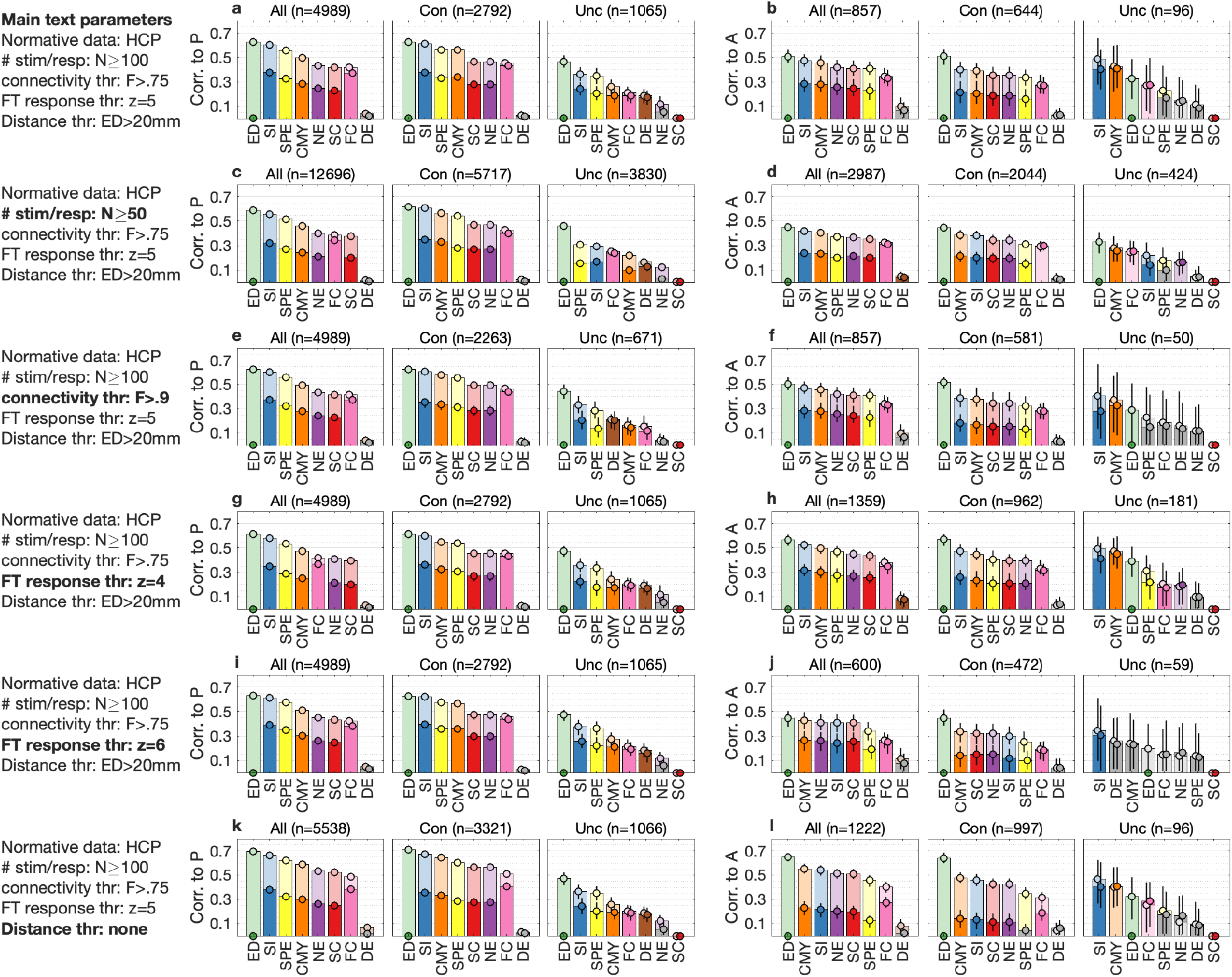
Control analyses for normative brain connectivity inferred from the Human Connectome Project (HCP) dataset. Each row of plots shows results obtained for a different set of methodological choices. To facilitate the comparison of control analyses and main text results, panels **(a–b)** reproduce the results reported in Fig 3. The main text results were obtained considering the following parameters: (i) pairs of regions were included in the analyses of response probability (amplitude) if they had at least 100 stimulation experiments (significant responses); (ii) pairs of regions were considered to be anatomically connected (unconnected) if they shared a connection in at least 75% (no more than 25%) of participants; (iii) response to stimuli were considered significant for *z*-scored CCEP amplitude larger than 5; and (iv) pairs of regions were included if they were at least 20mm apart in Euclidean distance. **(c-d)** Control analysis considering region pairs with at least 50 stimulation experiments (probability) or significant responses (amplitude). **(e-f)** Control analysis in which pairs of regions were considered to be anatomically connected (unconnected) if they shared a connection in at least 90% (no more than 10%) of participants. **(g–h)** Control analysis in which response to stimuli were considered significant for *z*-scored CCEP amplitude larger than 4. **(i–j)** Control analysis in which response to stimuli were considered significant for *z*-scored CCEP amplitude larger than 6. **(k–l)** Control analysis in which no restrictions were imposed on Euclidean distance between pairs of regions.

**FIG. S3.**
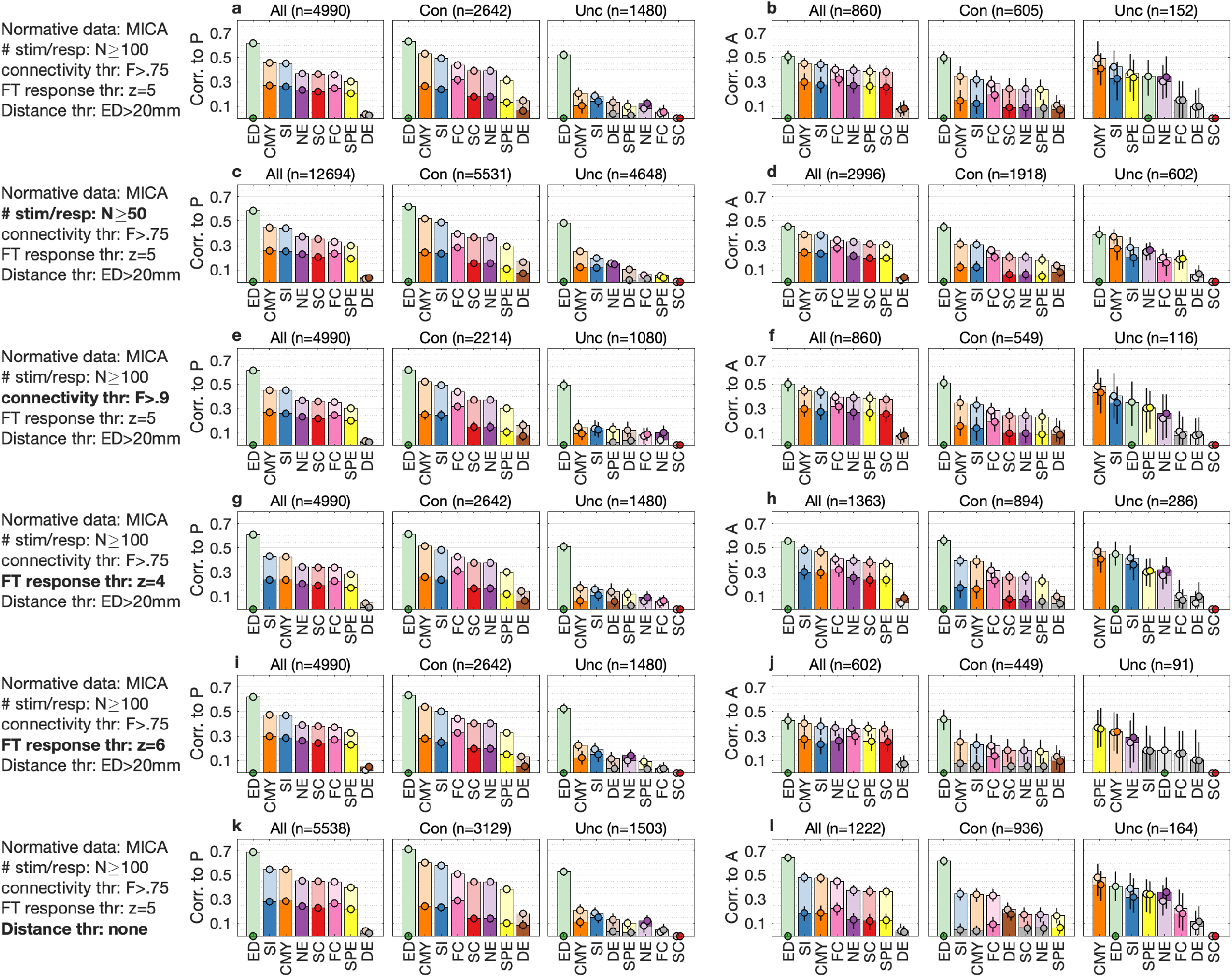
Control analyses for normative brain connectivity inferred from the Multimodal and Connectome Analysis (MICA) dataset. Each row of plots shows results obtained for a different set of methodological choices. Panels **(a–b)** shows results obtained considering the parameters utilized in the main text: (i) pairs of regions were included in the analyses of response probability (amplitude) if they had at least 100 stimulation experiments (significant responses); (ii) pairs of regions were considered to be anatomically connected (unconnected) if they shared a connection in at least 75% (no more than 25%) of participants; (iii) response to stimuli were considered significant for *z*-scored CCEP amplitude larger than 5; and (iv) pairs of regions were included if they were at least 20mm apart in Euclidean distance. **(c-d)** Control analysis considering region pairs with at least 50 stimulation experiments (probability) or significant responses (amplitude). **(e-f)** Control analysis in which pairs of regions were considered to be anatomically connected (unconnected) if they shared a connection in at least 90% (no more than 10%) of participants. **(g–h)** Control analysis in which response to stimuli were considered significant for *z*-scored CCEP amplitude larger than 4. **(i–j)** Control analysis in which response to stimuli were considered significant for *z*-scored CCEP amplitude larger than 6. **(k–l)** Control analysis in which no restrictions were imposed on Euclidean distance between pairs of regions.

**FIG. S4.**
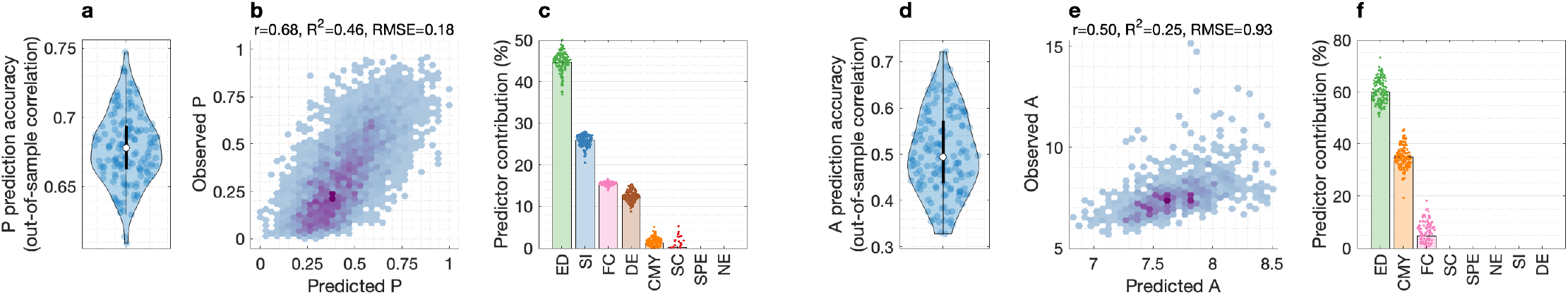
Machine learning predictions of stimulus response using the Multimodal and Connectome Analysis (MICA) dataset. Normative brain connectivity measures were independently inferred from different healthy adults, scanners, acquisition protocols, pre-processing techniques, and connectivity mapping methods. **(a)** Out-of-sample prediction accuracy for 15 repetitions of 10-fold cross validation, computed as the Pearson correlation between predicted and observed probabilities. **(b)** Scatter plot of predicted versus observed probabilities with prediction accuracy evaluated as the Pearson correlation, *R*^2^, and root mean squared error (RMSE). **(c)** Average contribution of input predictors across 15 repetitions of 10-fold cross validation. **(d–f)** Same as (a–c) but for predictions of response amplitude.

**FIG. S5.**
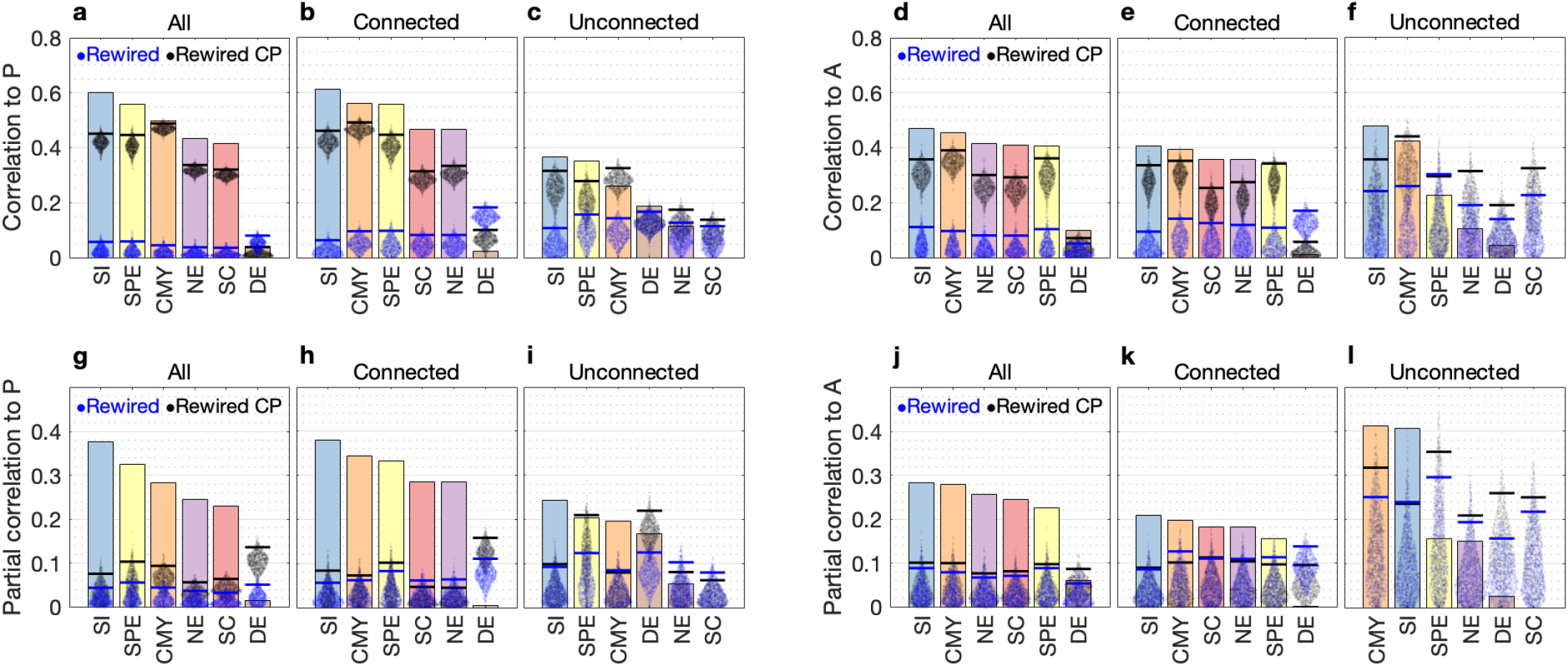
Comparisons of results obtained for network communication measures computed on empirical and topologically randomized SC. Bars show the empirical Spearman correlations reported in Fig 3. Point clouds show the distributions of null correlations obtained for 1,000 randomizations of SC using standard (blue) and cost-preserving (black) implementations of the Maslov-Sneppen rewiring algorithm. An empirical correlation outperforms the null models if the height of the bar exceeds the 95th percentile of null correlations marked by horizontal dashes (non-parametric significance test at *α* = 5%). Results for full Spearman correlations with response probability for **(a)** all, **(b)** anatomically connected, and **(c)** anatomically unconnected region pairs. **(d–e)** Same as (a–c) but for response amplitude. **(g–l)** Same as (a–f) but for partial Spearman correlations controlling for the effect of Euclidean distance.

